# Psychophysiological Interactions in a Visual Checkerboard Task: Reproducibility, Reliability, and the Effects of Deconvolution

**DOI:** 10.1101/133454

**Authors:** Xin Di, Bharat B. Biswal

## Abstract

Psychophysiological interaction (PPI) is a regression based method to study task modulated brain connectivity. Despite its popularity in functional MRI (fMRI) studies, its reliability and reproducibility have not been evaluated. We investigated reproducibility and reliability of PPI effects during a simple visual task, and examined the effect of deconvolution on the PPI results. A large open-access dataset was analyzed (n = 138), where a visual task was scanned twice with repetition times (TRs) of 645 ms and 1400 ms, respectively. We first replicated our previous results by using the left and right middle occipital gyrus as seeds. Then ROI-wise (regions of interest) analysis was performed among twenty visual-related thalamic and cortical regions, and negative PPI effects were found between many ROIs with the posterior fusiform gyrus as a hub region. Both the seed-based and ROI-wise results were similar between the two runs and between the two PPI methods with and without deconvolution. The non-deconvolution method and the short TR run in general had larger effect sizes and greater extents. However, the deconvolution method performed worse in the 645 ms TR run than the 1400 ms TR run in the voxel-wise analysis. Given the general similar results between the two methods and the uncertainty of deconvolution, we suggest that deconvolution may be not necessary for PPI analysis on block-designed data. Lastly, intraclass correlations between the two runs were much lower for the PPI effects than the activation main effects, which raise cautions on performing inter-subject correlations and group comparisons on PPI effects.

## 1. Introduction

Psychophysiological interaction (PPI) is a widely used method to study task related brain functional connectivity changes (Friston et al., 1997). It employed simple regression-based method to model task modulated connectivity effects, thus enabling whole brain exploratory analysis. Therefore, even though there are more sophisticated methods available, e.g. dynamic causal modeling (Friston et al., 2003), PPI is still a valuable method for fMRI data, given that our knowledge on large-scale task related connectivity is still quite limited. Several modifications of the PPI method have been made after it was proposed, including adding a deconvolution step to deal with the asynchrony between task design and fMRI hemodynamic response (Gitelman et al., 2003) and introducing a generalized framework to model more than two experimental conditions (McLaren et al., 2012).

A PPI effect is defined as an interaction between the time series of a brain region (physiological variable) and a (or more) task design variable (psychological variable). Noises of both the physiological and psychological variables go into the interaction term, so that the interaction effect is much noisier than the main effects of task free connectivity (physiological main effect) and task activation (psychological main effect). This makes PPI analysis having lower statistical power than simple connectivity and conventional activation analysis. Since PPI analysis has been increasingly used to study group differences and inter-subjects variability, it is important to evaluate the reproducibility and reliability of the PPI methods (Dubois and Adolphs, 2016; Vul et al., 2009). Voxel-based meta-analysis has been used to examine consistency of PPI results across studies (Di et al., 2017a). However, because the tasks used in different studies varied greatly, the motivation of a meta-analysis on PPI was rather to identify different connectivity that were modulated by different tasks, than to simply identify consistent connectivity cross studies with different tasks (Di et al., 2017a). Nevertheless, the reliability of PPI effect has not been directly examined.

One critical step for the PPI method is to properly deal with the asynchrony between task design and observed blood-oxygen-level dependent (BOLD) signals. An earlier solution is to convolve the psychological variable with hemodynamic response function (HRF). Then the PPI term *x*^1^_*PPI*_ could be expressed as:

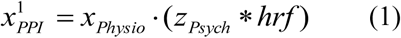

where *xPhysio* represents the physiological variable, *zPsych* represents the psychological design variable, and * represents convolution operator. However, this calculation is not appropriate if the interaction happened faster than the slow hemodynamic response. Therefore, a deconvolution procedure is required (Gitelman et al., 2003) to find a variable *zPhysio* that:

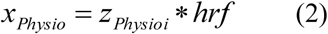

If this could be achieved, then the interaction could be calculated at the neuronal level and then convolve with HRF:

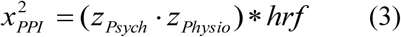

We can also put equation (2) to equation (1), so that:

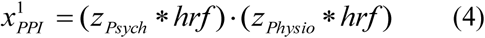

Mathematically, 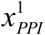 and 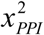 are not equivalent. Therefore, deconvolution seems necessary. Effective deconvolution depends on assumptions such as known HRF and noise characteristics in the BOLD signals (O’Reilly et al., 2012; Roebroeck et al., 2011). Unfortunately, there are substantial amount of variability in HRF both across brain regions and across subjects (Handwerker et al., 2004). On the other hand, if a task design is slower than the hemodynamic response, e.g. a blocked design, the PPI terms calculated from the above mentioned two methods could be very similar. We have demonstrated that the PPI results of a block-designed visual task are spatially corresponding very well between the deconvolution and non-deconvolution PPI methods (Di et al., 2017b). Whether to perform deconvolution then needs to compromise between the deviation between the PPI terms calculated in different ways and the uncertainty of deconvolution (Di et al., 2017b). Therefore, it might be better to not perform deconvolution for an block-designed task, which is actually recommended by FSL (FMRIB Software Library) (Jenkinson et al., 2012; O’Reilly et al., 2012). For event-related designed task, however, deconvolution may be still necessary, because the PPI terms calculated from the deconvolution and non-deconvolution methods may be dramatically different.

We recently demonstrated negative PPI effects (reduced connectivity) between the middle occipital gyrus to the fusiform gyrus and supplementary motor areas in a simple block-designed checkerboard task compared with a fixation baseline (Di et al., 2017b). Here, we further analyzed a larger sample of checkerboard data (n = 138) of two separate runs with two repetition times (TR: 645 ms and 1400 ms) (Nooner et al., 2012). The aims of the current study are to first evaluate reproducibility and reliability of PPI effects in the checkerboard task. Additionally, we investigated the impact of PPI calculation methods on the PPI results and their reproducibility and reliability. We operationally defined reproducibility as whether previously reported clusters could be observed in the current analysis, and whether the clusters reported in one run could be observed in the other run. Quantitatively, we utilized Dice coefficient to quantify overlaps of voxels on thresholded maps (Rombouts et al., 1998; Taylor et al., 2012). Next, we used intraclass correlation (ICC) to quantify test-retest reliability. Because the short TR run has about twice the number of time points as the long TR one, we predict that statistical results would be better for the short TR run compared with the long TR run. In addition, shorter sampling rate may provide more accurate estimate of hemodynamic response, therefore deconvolution PPI method should work better for the short TR than the long TR runs.

## 2. Methods

### 2.1. Simulations on the correlations between PPI terms

The hemodynamic response is a slow response compared with neuronal events, which can be understood as a low-pass filter. Intuitively, if a task design is slow enough, e.g. a blocked design, the convolution with the HRF may not affect PPI calculations much. To directly demonstrate this relationship between design alternating length and the effect of convolution on PPI calculation, we firstly performed a simulation. In this simulation, we defined a simple block-designed task with equal on and off periods with different cycle lengths (from 8 s to 80 s), and a simple event-related design with fixed inter trial interval of 12 s (Figure 1A). We used a typical sampling rate of 2 s, so that the event-related design could be expressed as alterations of one time bin (2 s) of a trial and 5 time bins (10 s) of the baseline condition (The first column in Figure 1A). The remaining columns in Figure 1A show block designs with different frequencies of repetition. For example, 80 secs cycle means 40-s on and 40-s off of the task condition related to the baseline. We then simulated the physiological variable of neuronal activities as a Gaussian variable for 1,000 times. For each design and simulated “neuronal” physiological variable, we calculated PPI terms using two ways: 1) each variable convolved with the canonical HRF and then the two convolved variables were multiplied to form a PPI term (corresponding to 1 *PPIx* in equation 4); 2) the two variables were multiplied and then convolved with the canonical HRF (corresponding to 2 *PPI x* in equation 3). We then calculated the correlations of the PPI terms calculated from the two methods. The code for this simulation can be found at: https://github.com/dixy0/PPI_correlation_demo.

**Figure 1.**
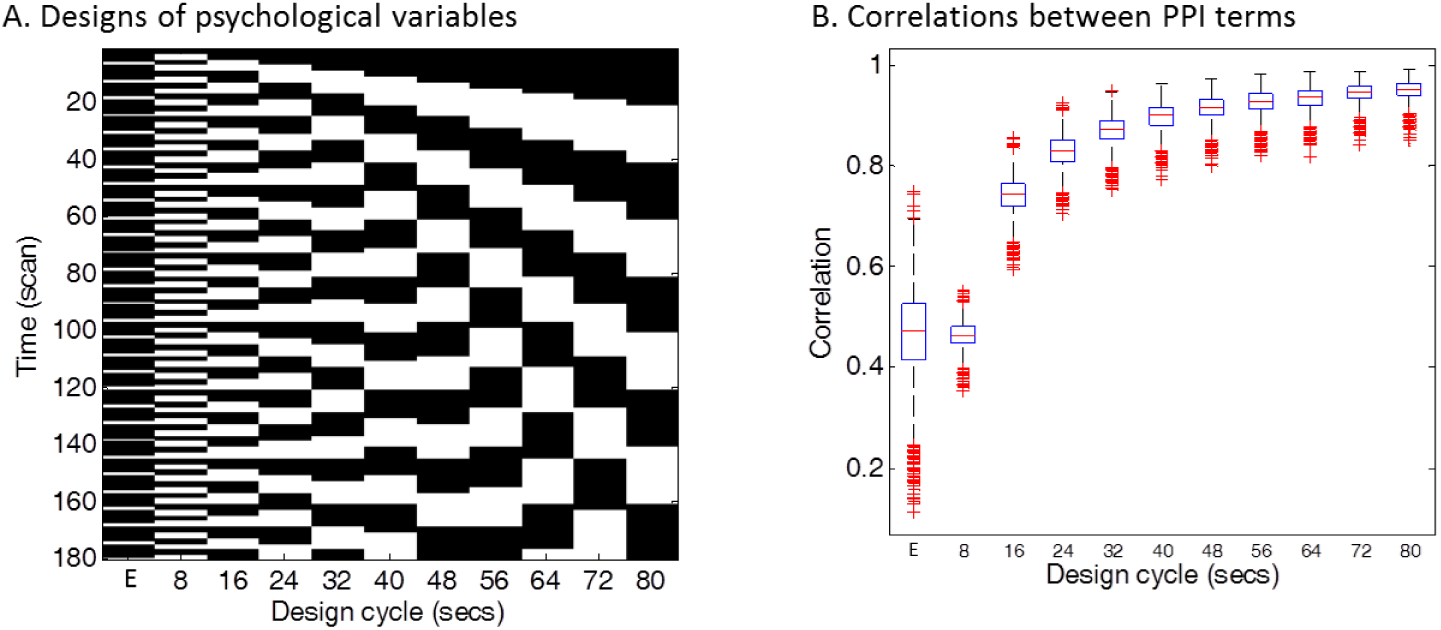
Simulations of the correlations between PPI terms calculated from deconvolution and non-deconvolution methods. Panel A illustrates different task designs that were used for the simulation. Each column represents a task design. E in the x axis represents the event-related design, with 1 time bin (2 s) of trial condition and 5 time bins (10 s) of baseline condition. The remaining columns show block designs with different frequencies of repetition. For example, 80 secs cycle means 40-s on and 40-s off of the task condition related to the baseline. Physiological variables at the neuronal level were generated using Gaussian random variables for 1,000 times. Penal B shows boxplots of correlations across the 1,000 simulations between PPI terms calculated from two methods: 1) the two simulated variables were convolved with the HRF and then multiplied to form the PPI term; 2) the two simulated variables were multiplied and then convolved with the HRF.

### 2.2. FMRI data and task design

We used the checkerboard fMRI data with TRs of 645 ms and 1400 ms from the release 1 of Enhanced Nathan Kline Institute - Rockland Sample (http://fcon_1000.projects.nitrc.org/indi/enhanced/). 146 subjects data with age equal or larger than 20 years old were included for analysis. Six subjects data were discarded due to large head motion during fMRI scanning in any of the two scans (maximum frame-wise displacement (FD) (Di and Biswal, 2015) greater than 1.5 mm or 1.5^o^). One subject s data were deleted because of poor coverage of the lower occipital lobe, and another subject s data were deleted because of failure of coregistration and normalization. The effective number of subjects was 138 (89 females, 45 males, 1 unidentified). The mean age of the sample was 47.8 years (20 to 83 years).

The checkerboard task consisted of 20 s fixation block and 20 s flickering checkerboard block repeated three times. A blank screen was presented after the third checkerboard block until fMRI scan was complete. The task was scanned for two separate runs with two TRs: 645 ms and 1400 ms, respectively. For the 645 ms run, 239 or 240 fMRI images were scanned for each subject. The following parameters were used: TR = 645 ms; TE = 30 ms; flip angle = 60 deg; voxel size = 3 × 3 × 3 mm^3^ isotropic; number of slices = 40. For the 1400 ms run, 98 fMRI images were scanned for each subject. The following parameters were used: TR = 1400 ms; TE = 30 ms; flip angle = 65 deg; voxel size = 2 × 2 × 2 mm^3^ isotropic; number of slices = 64. Anatomical T1 images were scanned using MPRAGE (magnetization-prepared rapid acquisition with gradient echo) sequence with the following parameters: TR = 1900 ms; TE = 2.52 ms; flip angle = 9°; voxel size = 1 × 1 × 1 mm^3^ isotropic. More information of the data can be found in Nooner et al. (Nooner et al., 2012).

### 2.3. FMRI data analysis

#### 2.3.1. FMRI data preprocessing

Functional MRI data preprocessing and analysis were performed using SPM12 software (http://www.fil.ion.ucl.ac.uk/spm/) under MATLAB environment (http://www.mathworks.com/). For the 645 ms run, the first 14 images (9 s) were discarded from analysis, resulting in 225 images for each subject. For the 1400 TR run, the first five images (7 s) were discarded from analysis, resulting in 93 images for each subject. The functional images were motion corrected, and corregistered to subject’s anatomical images. The anatomical images were segmented, and the deformation field images were used to normalize the functional images into MNI space. The data from the two TR runs were both resliced and resampled at a spatial resolution of 3 × 3 × 3 mm^3^. Lastly, the functional images were smoothed using a 6 mm full width at half maximum (FWHM) Gaussian kernel.

#### 2.3.2. Activation analysis

We first defined functional ROIs of the visual thalamus and lower visual area by performing general linear model (GLM) analysis on the checkerboard task. The checkerboard task was modeled as a box-car function, with 1 representing the checkerboard condition and 0 representing the fixation or blank screen. The box-car function was convolved with the canonical hemodynamic response function (HRF) to form a predictor of BOLD responses. Two regressors of the first eigenvariate of BOLD signals in white matter and cerebrospinal fluid (CSF), and 24 regressors of Friston s autoregressive head motion model (Friston et al., 1996) were also added in the model as covariates. An implicit high-pass filter of 1/128 Hz was also implemented in the model. The high-pass filtering is accomplished in SPM by using discrete cosine transform functions. The effective high-pass filtering cutoffs were then 0.0069 Hz (1 / 145.125 s) for the 645 ms TR run and 0.0077 Hz (1 / 130.2 s) for the 1400 ms TR run. The GLM model was estimated for each voxel in the brain to identify regions that showed similar patterns of activations as the task design. The beta maps of task activation were used for group level analysis using a one sample t-test model. Statistical significant clusters were identified by using cluster level statistics based on random field theory. Clusters were first identified using a one-tailed t-test at p < 0.001, and cluster extent was determined using false discovery rate (FDR) at p < 0.05.

#### 2.3.3. Definition of regions of interest

We performed two types of PPI analyses, voxel-wise analysis using seed regions that were activated by the checkerboard task and ROI-based analysis among visual thalamus and cortical visual areas independently defined from other toolbox. In the activation analysis of the current data, the posterior visual cortex and the posterior portion of the thalamus were robustly activated by the visual checkerboard stimulation in both TR runs. We therefore defined the left and right middle occipital gyrus (LMOG and RMOG) and the thalamus as regions of interest (ROIs) based on the activations. To define the ROIs with proper size, we increase the threshold to t > 16 to define the LMOG and RMOG, and made an intersection between the two runs. The size of LMOG was 222 voxels, and the size of RMOG was 259 voxels. Thalamus was defined using a threshold of p < 0.001, with an intersection between the two runs. Because the visual thalamus is small, left and right ROIs were combined to form a single thalamus ROI (171 voxels). Different thresholds were chosen to ensure that these ROIs are similar in size. The eigenvariate of a ROI was extracted with adjustment of effects of no interests (head motion, WM/CSF variables, and low frequency drifts).

We defined the visual thalamus as the regions that show functional associations with the lateral visual network in resting-state (Yuan et al., 2016). Cortical visual areas were defined by using probabilistic cytoarchitectonic maps. These areas include the OC1/OC2 (occipital cortex) (Amunts et al., 2000), ventral and dorsal OC3 and OC4 (Kujovic et al., 2013; Rottschy et al., 2007), OC5 (Malikovic et al., 2006), and FG1/FG2 (fusiform gyrus) (Caspers et al., 2013). For the probabilistic maps of these regions, we first performed a winner-takes-all algorithm to define unique regions of each area, and then split them into left and right regions. As a result, there are 20 ROIs (left and right thalamus, OC1, OC3, OC3d, OC3v, OC4d, OC4v, OC5, FG1, and FG2). The eigenvariate of a ROI was extracted with adjustment of effects of no interests (head motion, WM/CSF variables, and low frequency drifts).

#### 2.3.4. Psychophysiological interaction analysis

PPI analysis was performed using SPM12 with updates 6685. PPI terms were calculated by using both deconvolution method and non-deconvolution method. For the deconvolution method, the time series of a seed region was deconvolved with the canonical HRF, multiplied with the centered psychological boxcar function, and convolved back with the HRF to form a predicted PPI time series at hemodynamic response level. For the non-deconvolution method, the box-car function of psychological design was convolved with the HRF to form a psychological variable, and it was centered and multiplied with the raw seed time series. Figure 2 shows examples of PPI terms calculated from the two methods in the two TR runs.

**Figure 2.**
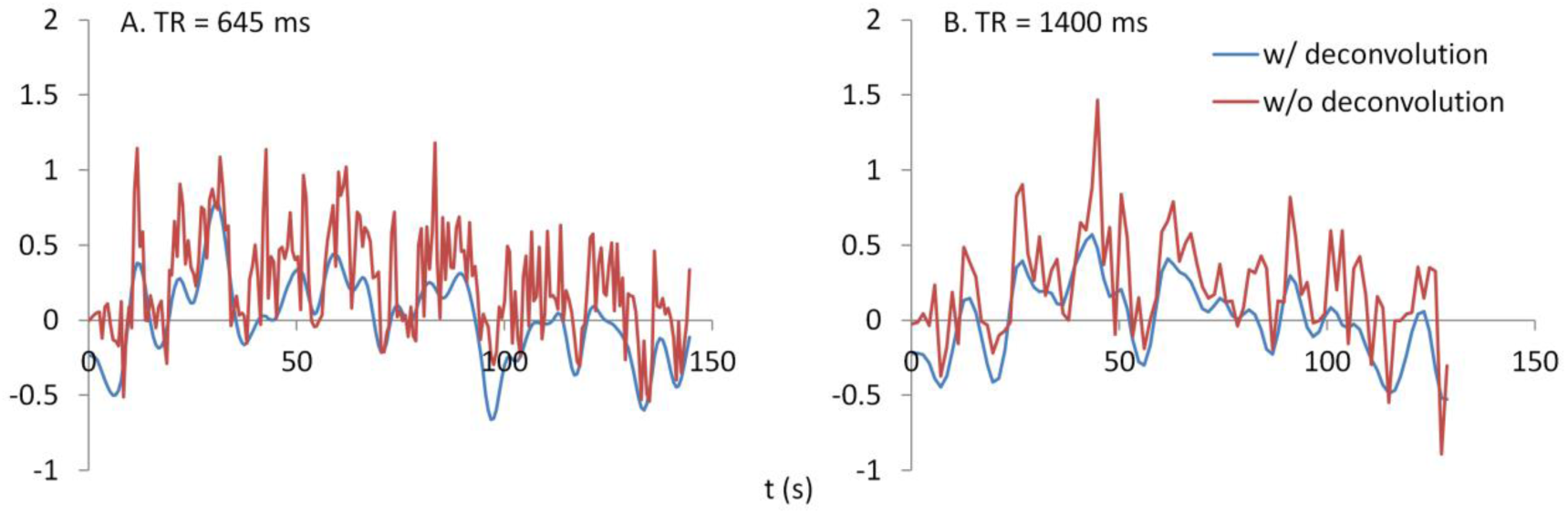
Examples of PPI terms calculated by the deconvolution and non-deconvolution methods for the two TR runs.

For voxel-wise PPI analysis, separate GLMs were built for the LMOG, RMOG, and thalamus seeds, and for the two TR runs. The models included one regressor representing task activation, one regressor representing the seed time series, the PPI term, and covariates the same as the activation GLMs descripted above. Group-level one sample t-test was used on the corresponding PPI effects, to test where in the brain showed consistent PPI effects with a seed region. For both positive and negative contrasts, a one-tailed t-test of p < 0.001 was first used to define clusters, and then a FDR cluster threshold of p< 0.05 was used to identify statistical significant clusters. For ROI-wise analysis, PPI GLM models were built for each of the 20 ROIs, and applied to all other ROIs as a dependent variable. The GLM model included one psychological variable, one physiological variable, one PPI variable, and one constant term. The covariates were not included because they have already been regressed out from all ROI time series. PPI effects were calculated between each pair of ROIs, resulting in a 20 × 20 matrix of beta values for each subject. The matrices were symmetrized by averaging corresponding upper and lower diagonal elements (Di et al., 2017b), with a total of 190 (20 × 19 / 2) unique effects. Group-level one-sample t-test was performed on each element of the matrix. For both positive and negative contrasts, a one-tailed t-test of p < 0.001 was used to identify significant PPI effects. This threshold was chosen to match with voxel-wise analysis. We also used FDR correction on the total of 190 effects. And the results are similar to what using a p < 0.001 threshold. However, FDR depends on the distribution of all tested p values, making it difficult to compare between two runs. Therefore, we adopted p < 0.001 to report ROI-based PPI results.

#### 2.3.5. Reproducibility and reliability

We operationally define reproducibility as overlaps of supra-threshold clusters. Dice coefficient was used to quantify reproducibility (Rombouts et al., 1998). Two strategies were used to threshold the maps or matrix from the two TR runs. First, statistical t maps or t matrices from the two TR runs were thresholded using a common t value, ranging from 1.7 (approximately corresponds to p < 0.05) to 7. However, it is possible that the effect sizes in the two TR runs are systematically different, so that using a same t value could generate very different numbers of supra-threshold voxels or elements in the two runs. Therefore, we also thresholded t maps or t matrices based on the percentile of t values within a map or matrix. This could ensure that the numbers of supra-threshold voxels or elements are the same between the two TR runs. We operationally define reliability as test-retest reliability between the two TR runs, as quantified as ICC (Zuo et al., 2010a). Voxel-wise ICC maps or each ROI and ICC matrices across 20 ROIs were calculated between two TR runs for each PPI method. At each voxel or matrix element, ICC was calculated from a 138 (subject) by 2 (run) matrix by using a MATLAB function written by Zuo et al. (Zuo et al., 2010a). Because only voxels that have significant effects might show meaningful reliability, we displayed histograms of ICCs within significant voxels or elements with reference to those in the whole brain. For task activations, the significant voxels were determined using intersection of the two TR runs each thresholded at p < 0.001. For PPI effects of each ROI, the significant voxels were determined using intersection of the two TR runs and two methods each thresholded at p < 0.01. This slightly liberal threshold was chosen to ensure enough number of voxels survived in the conjunction of the four scenarios. The whole brain mask was determined as all voxels in the brain, including WM and CSF.

#### 2.3.6. Coefficient of variation

We calculated coefficient of variation to estimate measurement error of task activations and PPI effects. Coefficient of variation was calculated in ROIs that showed significant activation effects. Specifically, the LMOG, RMOG, and thalamus ROIs that were used as seed in the PPI analysis were used to represent activation effects. For the PPI results, we performed a conjunction analysis of the voxel-wise negative PPI effects across all the eight contrasts (2 PPI methods × 2 TR runs × 2 seeds) using a threshold of p < 0.01, and identified 27 ROIs that showed common negative PPI effects. Beta values of activations or PPI effects of these ROIs were extracted. Coefficient of variation was calculated based on the method assuming the variation is proportional to the mean (Bland and Altman, 1996). It measures within subject variations (across the two TR runs in the current case) relative to the mean effects of the two runs. Specifically, coefficient of variation was calculated based on a 138 (subject) × 2 (run) matrix. The beta values were first logarithmic transformed. Variation was then calculated for each subject, and a square root of mean variations across subjects was calculated. The resulting value was then transformed back using an exponential function, and subtracted by 1. The script for calculating coefficient of variation is available at: https://github.com/dixy0/PPI_correlation_demo. The resulting value represents the percentage of variation of a measure relative to the mean. Coefficients of variation were calculated on the LMOG, RMOG, and thalamus ROIs to reflect measurement errors of the task activations, and were calculated on the 27 ROIs and two ROI seeds to reflect measurement errors of the PPI effects.

## 3. Results

### 3.1. Simulations on the correlations between PPI terms

The distributions of PPI correlations for each task design are shown in Figure 1B. For the block designs, the PPI correlations are a function of block cycle length. With longer design cycle, e.g. greater than 40 s (20-s on and 20-s off), the correlations of PPI terms could be higher than 0.9. Practically, most of the block-designed fMRI experiments have longer block cycles than 20-s on and 20-s off. If the block alterations become faster, the correlation between PPI terms decreased. And for the event-related design, the mean PPI correlations were below 0.5 and with large variations. This simulation demonstrates that if a neuronal activity time series is known, using convolved time series to calculate PPI term (i.e. 1 *PPIx*) could be very similar to what calculated by first multiplying the two variables and then convolving (i.e. 2 *PPI x*) for typical block designed experiments. In real fMRI data, the “neuronal” physiological variable is not known, and has to be estimated by using deconvolution. Considering the similarities of the PPI terms and the caveats of deconvolution, PPI calculations without deconvolution may be a better choice for block designed experiments. On the other hand, the PPI correlations in the event-related design are much smaller (r < 0.5, meaning less than 25% of shared variance). So that deconvolution is still a necessary step for PPI analysis in event-related designed experiments.

### 3.2. Activations of the checkerboard task

Both TR runs showed highly significant activations in the visual cortex, as well as in the posterior portion of the thalamus (Figure 3A). The overlaps (Dice coefficients) of thresholded t maps between the two TR runs were as high as 0.7 (Figure 3B) at most of the shown t range percentile range. And Dice coefficients went down when only extremely activated voxels were thresholded. The visual cortex regions also showed high test-retest reliability (ICC greater than 0.7) (Figure 3C). However, the activations of the thalamus only showed small test-retest reliability around 0.2. The histograms of ICCs in the significant voxels and in the whole brain are shown on the right of Figure 3C.

**Figure 3.**
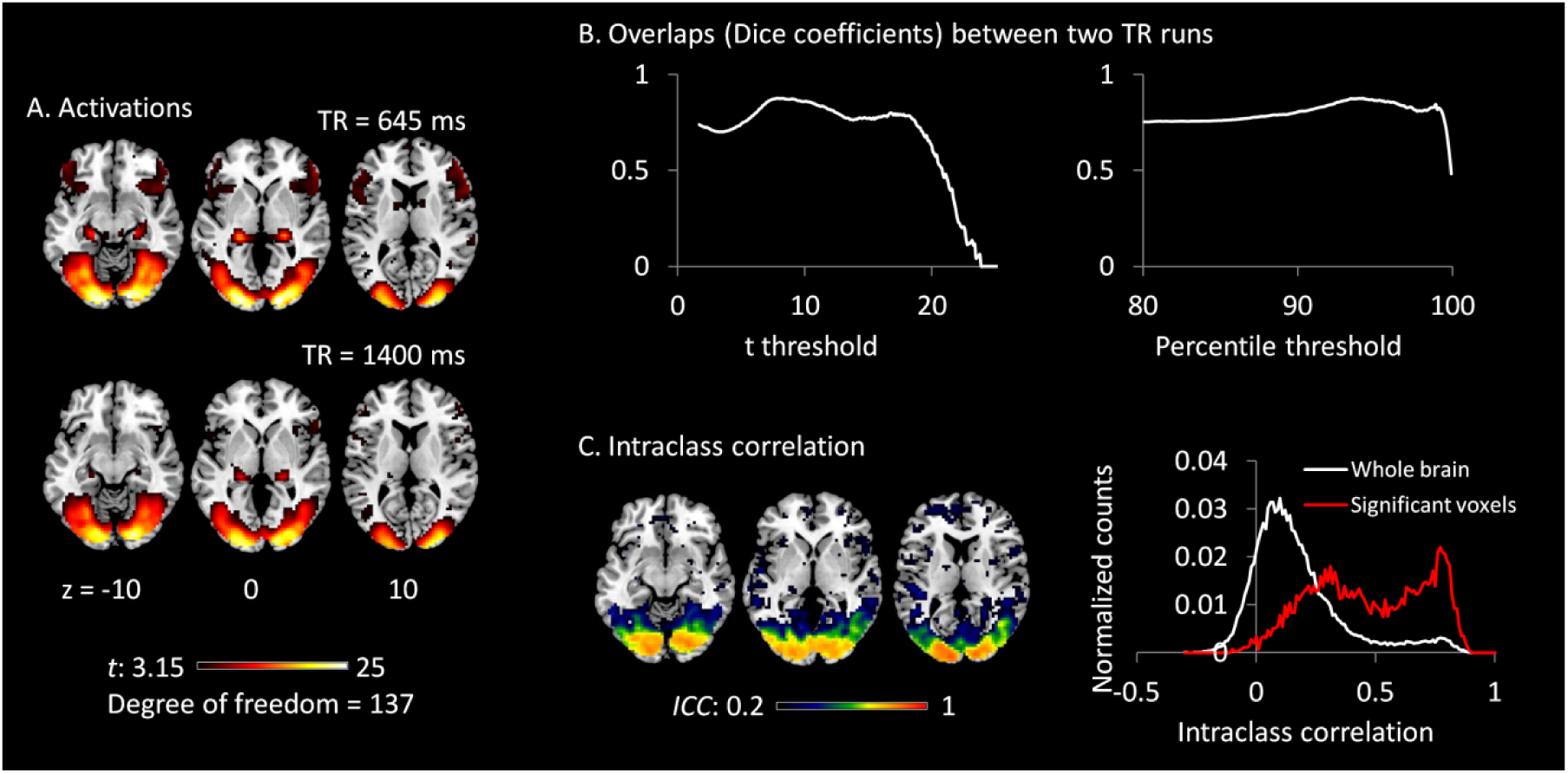
A) Activations (*t* maps) of visual checkerboard presentation for the 645 ms TR run (upper) and 1400 ms TR run (lower). The threshold t value corresponds to one-tailed significance at p < 0.001. B) Overlaps (Dice coefficients) between the two TR runs using t threshold (left) and percentile threshold (right). C) Test-retest reliability map (intraclass correlations, ICC) of activations between the two runs is shown on the left, which were thresholded at ICC > 0.2. The histograms of ICC of activations between the two TR runs in significant voxels and whole brain are shown on the right. The significant voxels were determined using intersection of the two runs each thresholded at p < 0.001.

### 3.3. Psychophysiological interactions

The voxel-wise PPI analysis of the LMOG and RMOG seeds conveyed very similar patterns. The PPI effects of the LMOG seed for the two TRs and two methods are shown in Figure 4. We first observed that even though spatial extents of PPI effects varied across the two TR runs and two PPI methods, the negative PPI effects in previously reported regions, i.e. supplementary motor area and higher visual cortex, could be observed from all four scenarios. The deconvolution method in 645 ms TR run had the smallest spatial extent and statistical significance, while the non-deconvolution method in 645 ms TR run had the largest spatial extent and strongest statistical significance. Both methods in TR of 1400 ms showed similar spatial extent and significance levels. The last row in Figure 4 demonstrates the overlaps of negative effects in the four scenarios. Similar results were found in the analysis of the RMOG seed (Figure 5).

**Figure 4.**
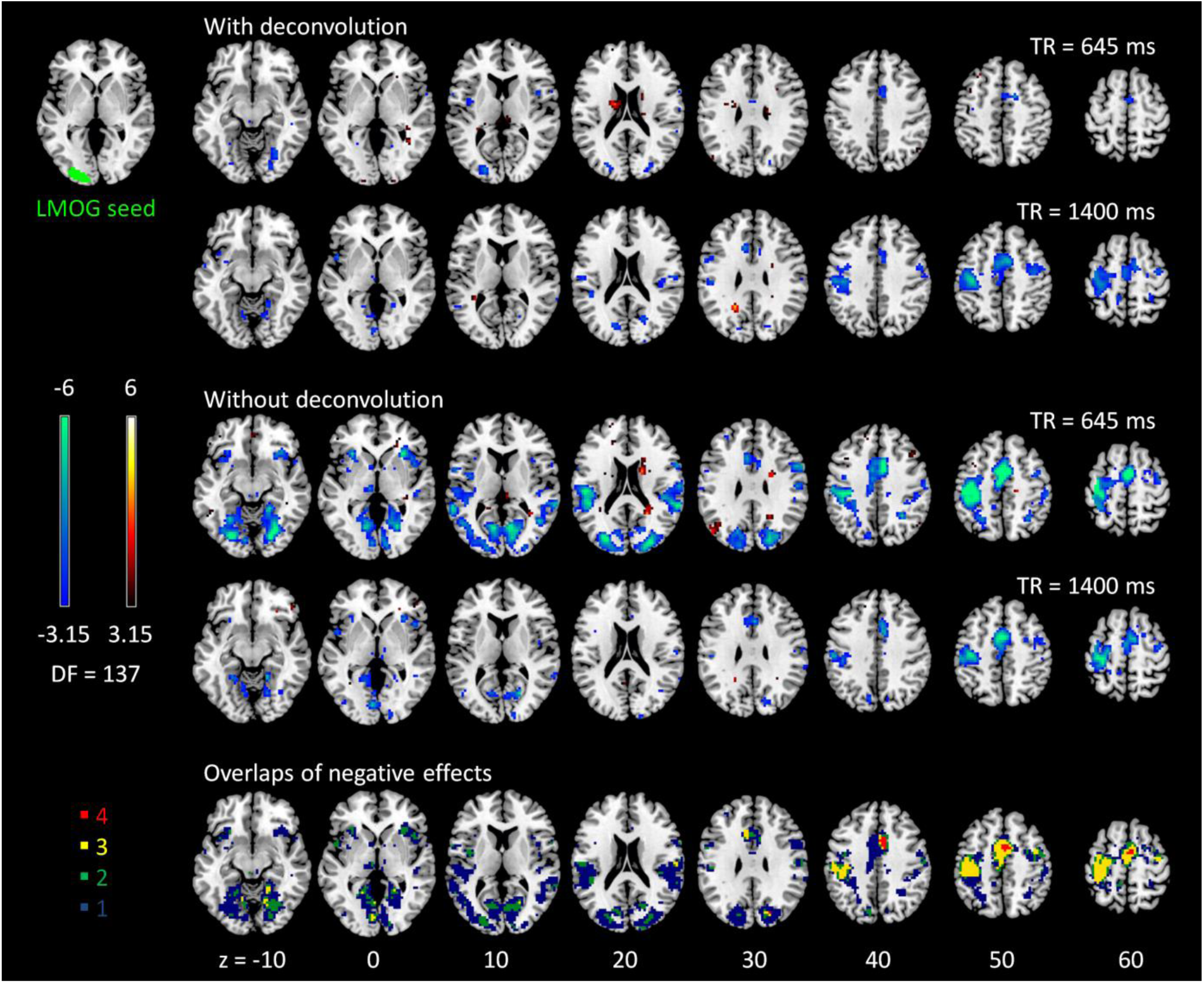
Psychophysiological interaction (PPI) results for the left middle occipital gyrus (LMOG) seed during checkerboard presentation in the two TR (repetition time) runs of 645 ms and TR 1400 ms. The resulting clusters were thresholded at p < 0.001 (approximated t = 3.15), with DF (degree of freedom) of 137. The last row illustrates the number of overlapped negative PPI results in the four scenarios. Numbers on the bottom represent z coordinates in MNI (Montreal Neurology Institute) space.

**Figure 5.**
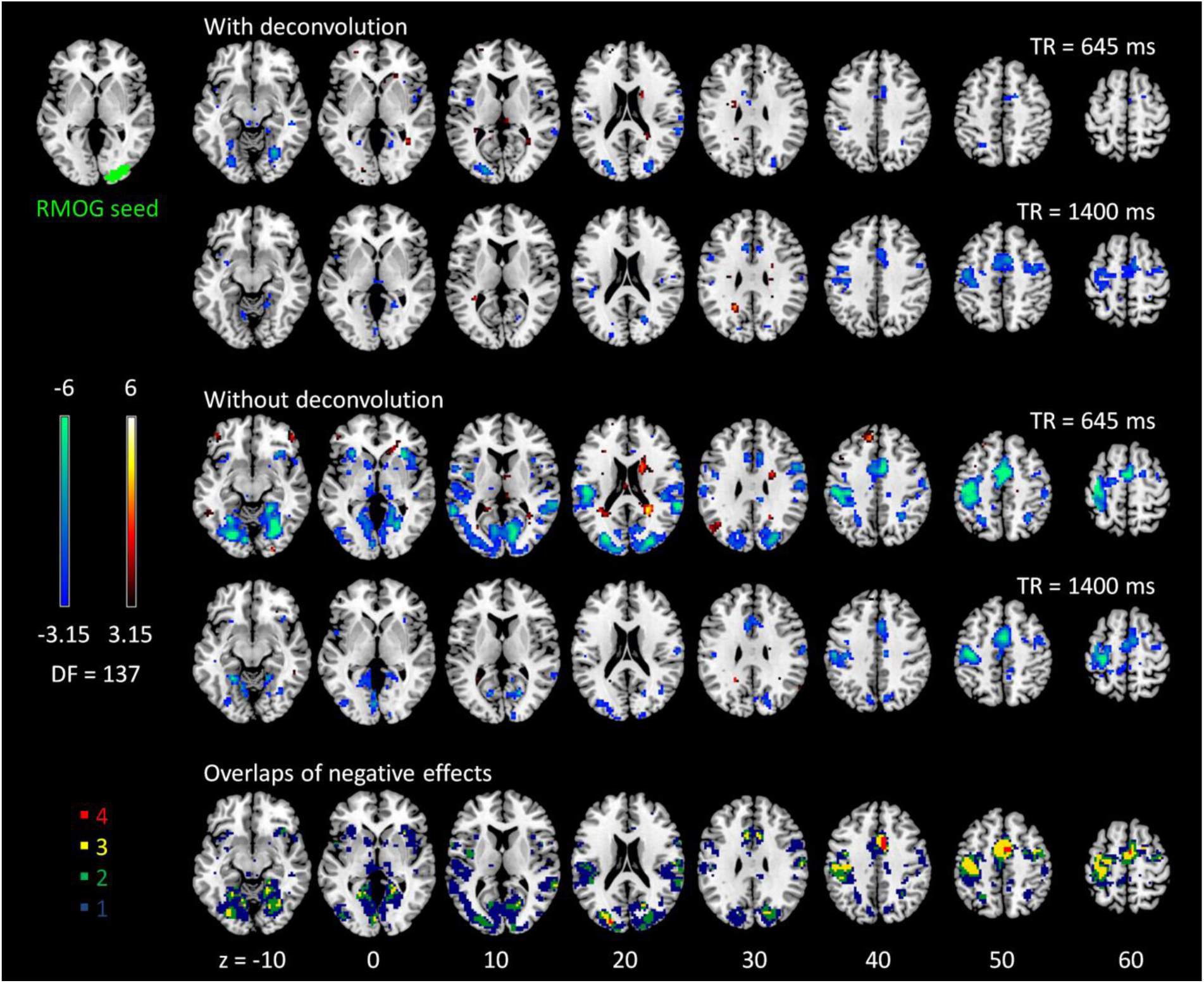
Psychophysiological interaction (PPI) results for the right middle occipital gyrus (RMOG) seed during checkerboard presentation in the two TR (repetition time) runs of 645 ms and TR 1400 ms. The resulting clusters were thresholded at p < 0.001 (approximated t = 3.15), with DF (degree of freedom) of 137. The last row illustrates the number of overlapped negative PPI results in the four scenarios. Numbers on the bottom represent z coordinates in MNI (Montreal Neurology Institute) space.

The voxel-wise PPI analysis of the thalamus seed only showed significant effects in the 645 TR run, but with different brain regions with opposite effects (Figure 6). With deconvolution method, the thalamus seed showed significant positive PPI effects with middle cingulate gyrus, anterior portion of the thalamus, bilateral anterior insula, basal ganglia, and right fursiform gyrus. Whereas with non-deconvolution method, the thalamus seed showed significant negative PPI effects with the bilateral occipital pole regions. There were no consistent results between two TR runs and two methods. Therefore subsequent analysis was only performed on the LMOG and RMOG seeds.

**Figure 6.**
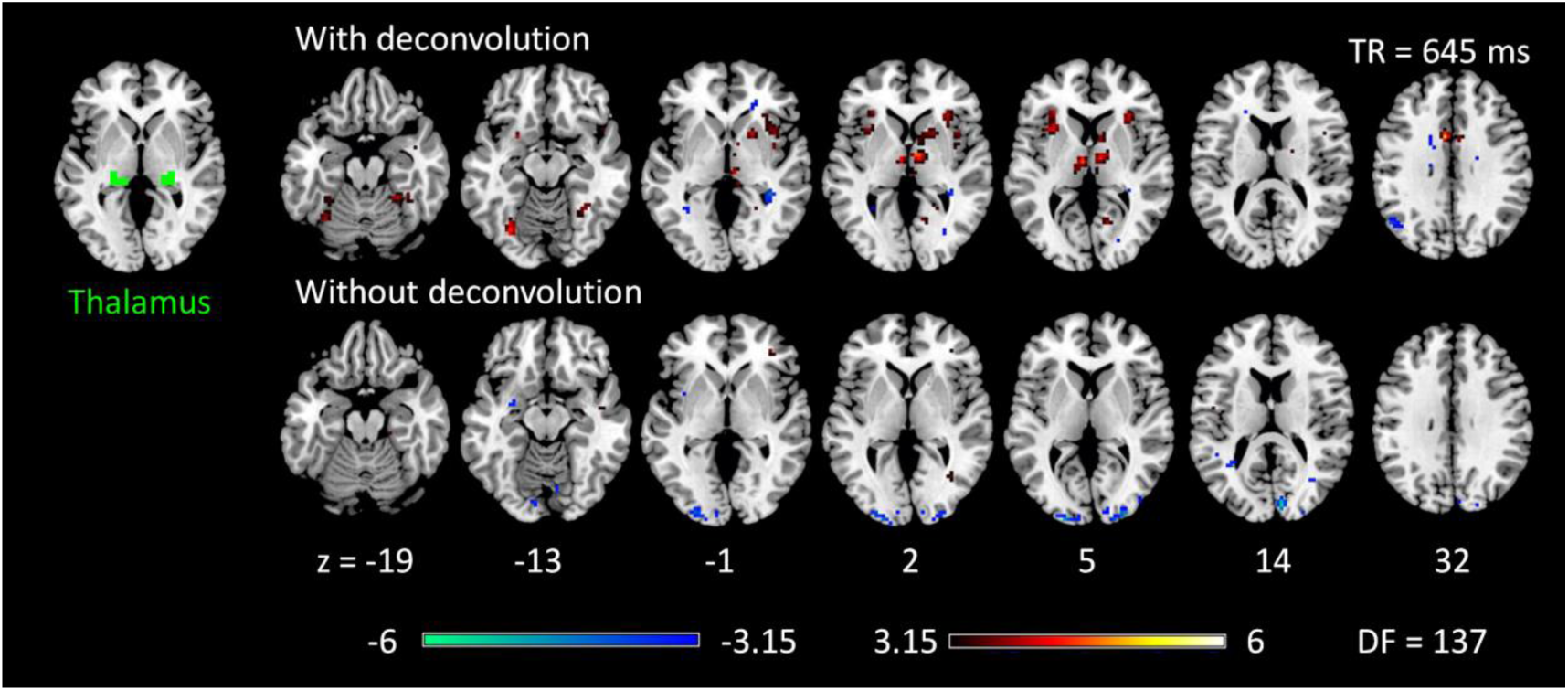
Psychophysiological interaction (PPI) results for the thalamus seed during checkerboard presentation in the TR (repetition time) run of 645 ms. There is no significant PPI effects of the thalamus seed in TR run of 1400 ms. The resulting clusters were thresholded at p < 0.001 (approximated t = 3.15), with DF (degree of freedom) of 137. Numbers on the bottom represent z coordinates in MNI (Montreal Neurology Institute) space.

We next performed ROI-based PPI analysis among 20 regions of visual thalamus and cortical visual areas (Figure 7). The 645 ms TR run showed more significant PPI effects than the 1400 ms TR run. And non-deconvolution method showed more significant PPI effects than the deconvolutio method. A prominent number of connectivity changes are between the bilateral FG1 regions and other lower level visual areas ranging from OC1, OC2, to OC4. We performed a conjunction analysis of PPI results across the four scenarios, and identified five connections with reduced connectivity in checkerboard than in fixation. The regions and connections are highlighted in Figure 8.

**Figure 7.**
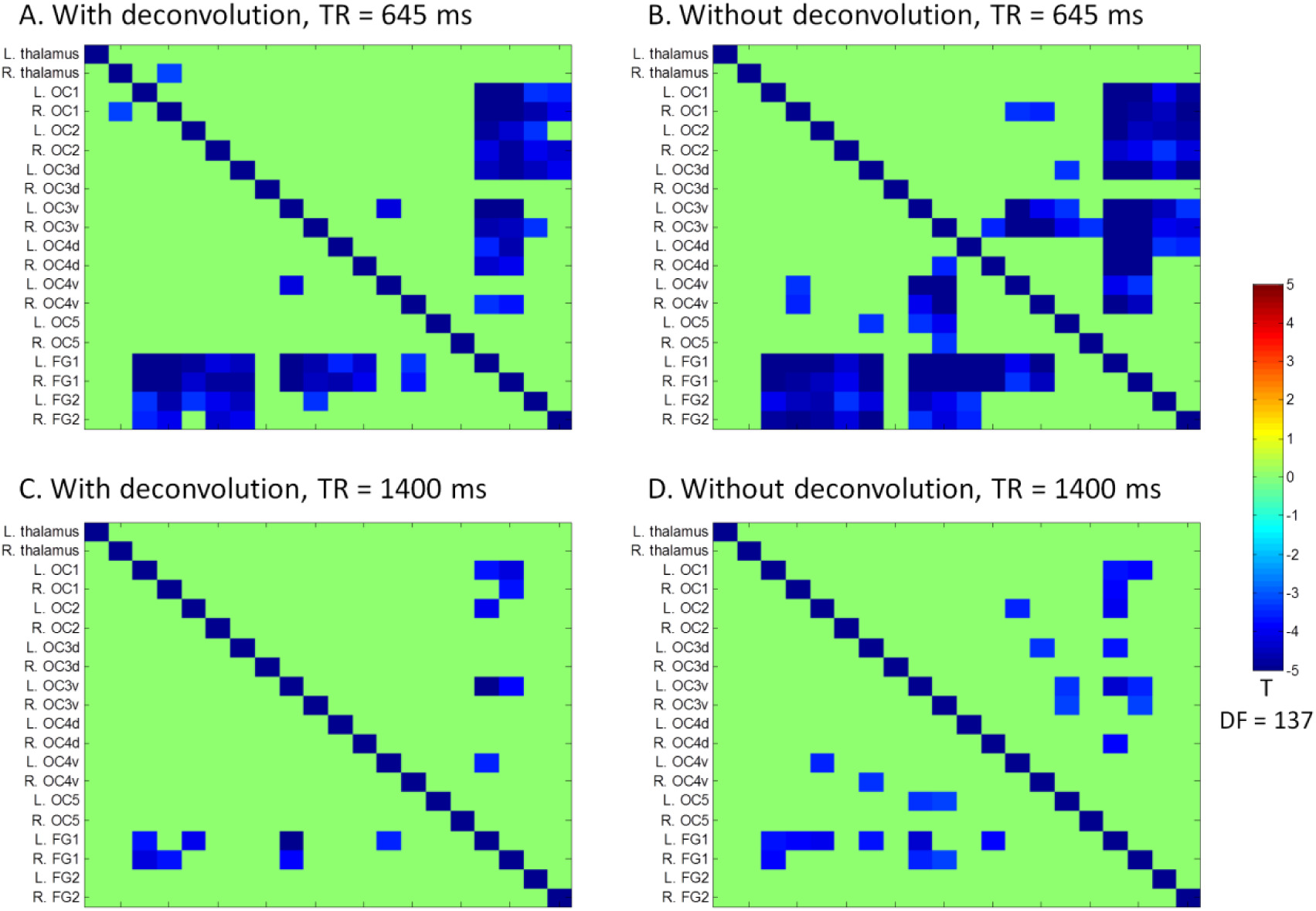
Matrices of psychophysiological interaction (PPI) results among the 20 regions of interest of visual thalamus and visual cortex for the two TR (repetition time) runs and two methods. The resulting clusters were thresholded at p < 0.001.

**Figure 8.**
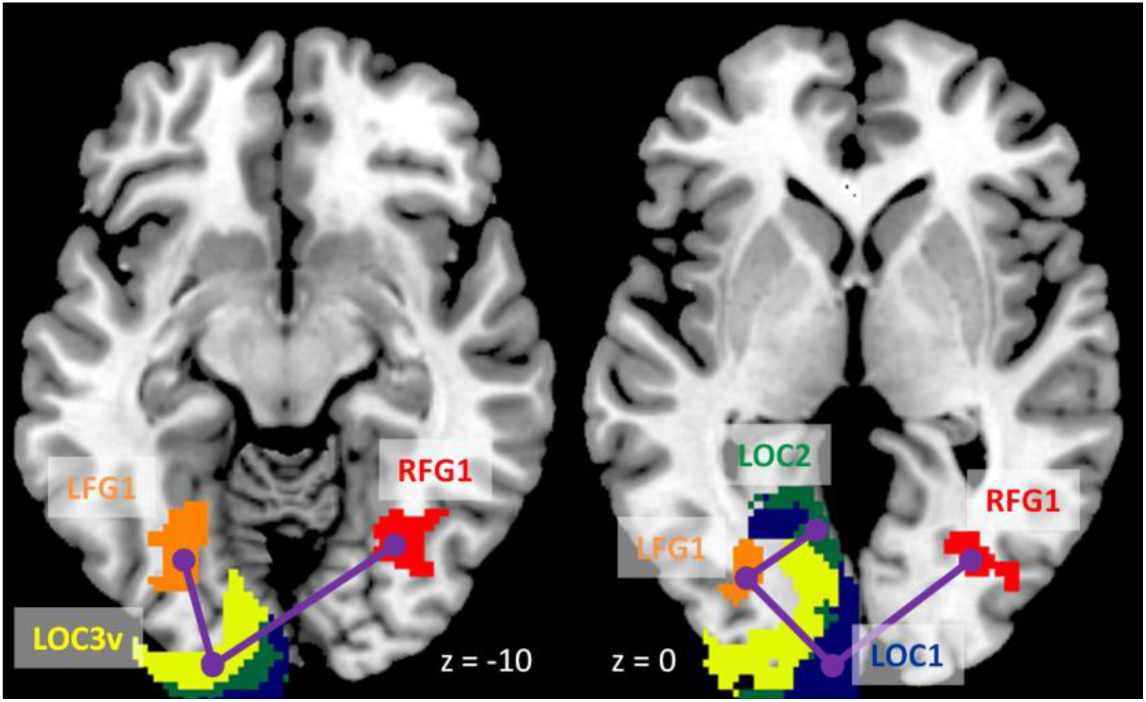
Illustration of consistently reduced connectivity during checkerboard presentation compared with fixation in the ROI-based (region of interest) psychophysiological interaction (PPI) analysis in the two TR (repetition time) runs and two methods. Numbers on the bottom represent z coordinates in MNI (Montreal Neurology Institute) space.

### 3.4. Reproducibility of PPI effects

Since we observed similarities of spatial clusters and connectivity between the two TR runs, we next examined reproducibility of PPI effects by calculating Dice coefficients of thresholded statistical maps or PPI matrices between the two TR runs (Figure 9). For voxel-wise analysis of both LMOG and RMOG seeds, when varying t threshold, the non-deconvolution method showed higher level overlap compared with the deconvolution method (Figure 9A). When thresholding statistical maps with matched number of surviving voxels, a similar pattern could still be observed that the non-deconvolution method produced larger overlaps than the deconvolution method (Figure 9B). For the ROI-wise analysis, however, Dice coefficients were at similar level between two PPI methods at most t and percentile thresholds. But at very high t threshold or percentile thresholds, the deconvolution method seemed to produce larger overlaps (higher Dice coefficients) (Figure 9C and 9D).

**Figure 9.**
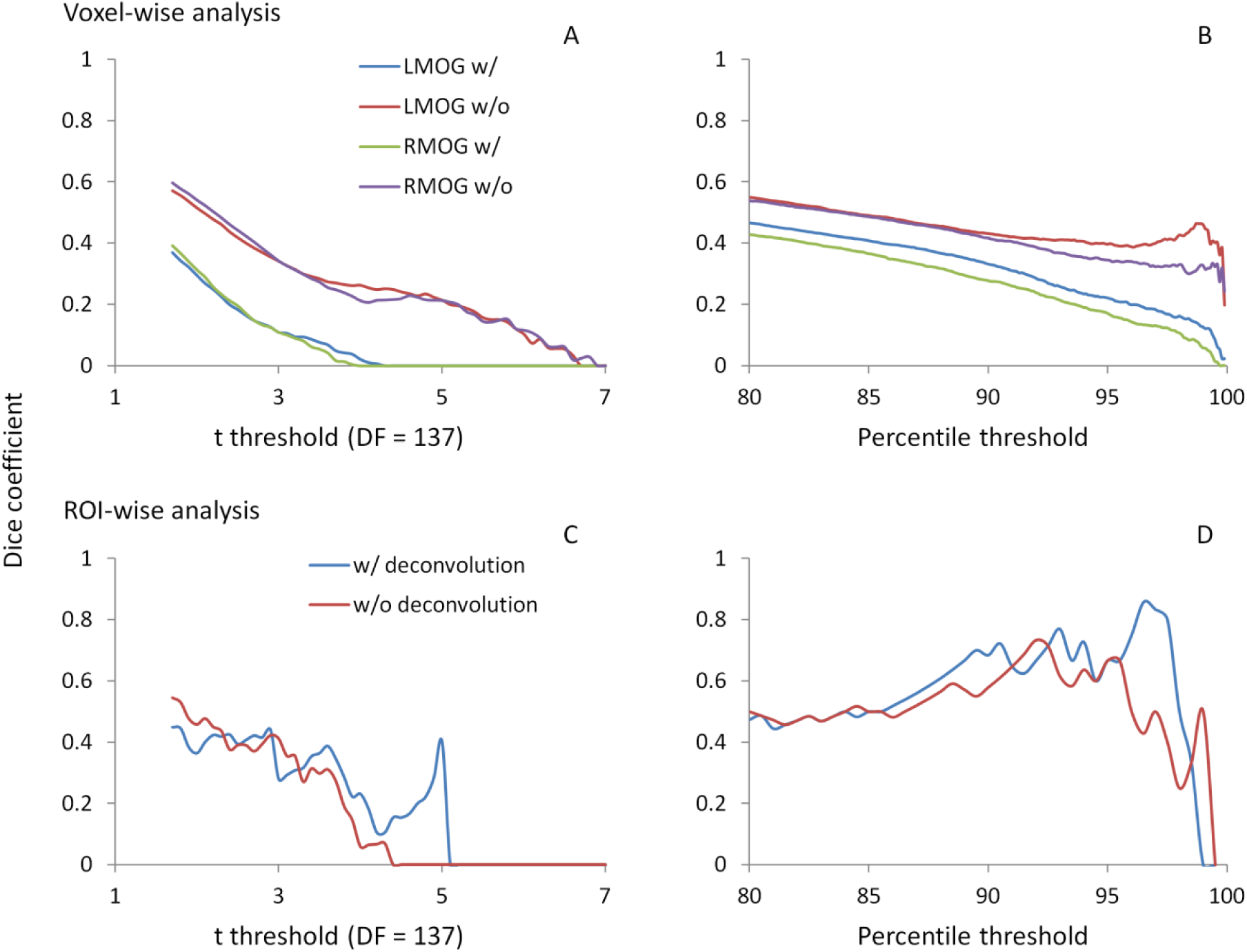
Dice coefficients of thresholded negative PPI effects between the two TR runs as functions of t threshold (A) and percentile threshold (B) for the two seeds and two PPI methods. The lowest t used for calculating overlap is 1.7, which approximately corresponds to p < 0.05. The largest percentile is 80 to 99.9 percentile, which is approximately corresponds to the largest proportions of voxels at p < 0.05.

### 3.5. Reliability of PPI effects

Lastly, we calculated ICC between the two TR runs to reflect reliability of PPI effects. The voxel-wise maps of ICC showed that there were typically low reliability in both methods and ROIs, even in the regions that showed consistent negative PPI effects (Supplementary Figure S1). We then plotted the histograms of ICCs in voxels from the whole brain (gray lines) and within regions that showed significant PPI effects (red lines) (Figure 10A through 10D). It turns out that the distributions of ICCs within significant regions are only slightly different from the distributions of correlations in the whole brain, with means around 0.07. The distributions of ICCs were not different between deconvolution and non-deconvolution methods. Similar distributions of ICCs were also found for the ROI-wise analysis (Figure 10E, 10F, and supplementary Figure S2). We found five PPI effects that were consistently significant in both TR runs and methods. And the ICCs for the five effects were also small and close to zero.

**Figure 10.**
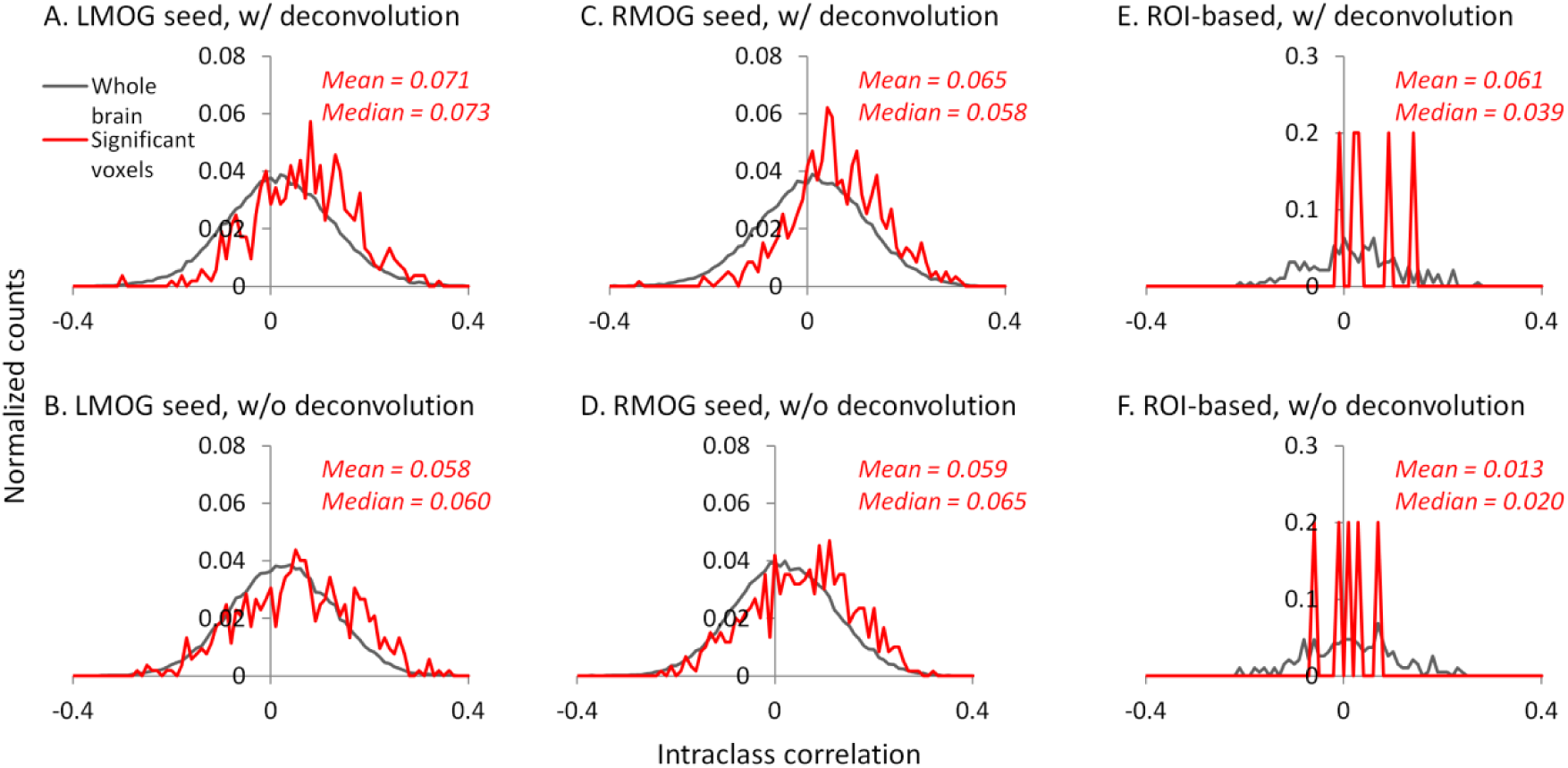
Histograms (normalized) of intraclass correlations of PPI effects between the two TR runs across the whole brain (gray lines) and in statistically significant voxels (red lines). The significant voxels were determined using intersection of the two runs and two methods each thresholded at p < 0.01. Left and right masks were calculated separately.

### 3.6. Measurement error

We calculated coefficients of variation (Bland and Altman, 1996) on task activations and PPI effects to reflect measurement error (Figure 11). The variations of activation in the LMOG and RMOG were about 70% of the mean activation, while the variation of activation in the thalamus was about 270% of the mean activation (Figure 11A). In contrast, the variations of PPI effects through the 27 ROIs were about 500% of the mean effects for both the LMOG and RMOG seeds (Figure 11 B and 11C), which indicated much arger variation of PPI effects compared with activations. The deconvolution and non-deconvolution methods had similar level of coefficients of variations. But when directly comparing the two methods, there was a trend that the non-deconvolution method had smaller coefficients of variation than the deconvolution method in most of the ROIs (Figure 11D and 11E).

**Figure 11.**
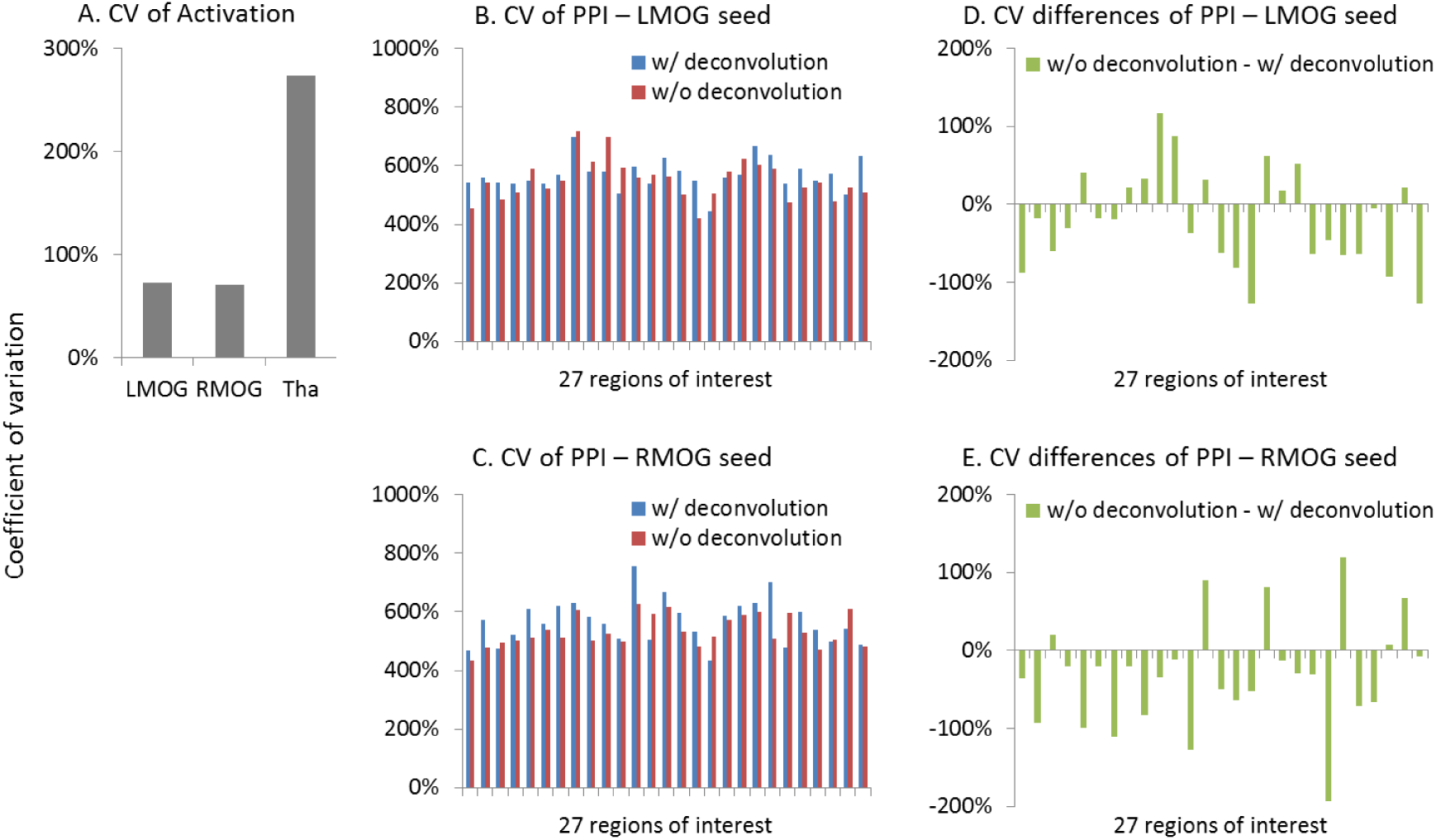
Measurement errors as revealed by coefficients of variations (CV) (Bland and Altman, 1996) for the activation (A) and psychophysiological interaction (PPI) results (B and C). Please notice the different scales in y axes. D) and E) demonstrate the differences of CV on PPI effects between the deconvolution and non-deconvolution methods. LMOG, left middle occipital gyrus; RMOG, right middle occipital gyrus; Tha, thalamus.

### 3.7. Miscellaneous analysis

To gain further insight to the cases of deconvolution failure, we calculated correlations of PPI terms between deconvolution and non-deconvolution methods for the LMOG and RMOG seeds (Figure 12A). In both TR runs, the distributions of correlations centered approximately on 0.7, and there were outliers whose correlations were only 0.2 or 0.3. This is in contrast with the simulation results (Figure 1B, 40 sec cycle), where the correlations were around 0.9.

**Figure 12.**
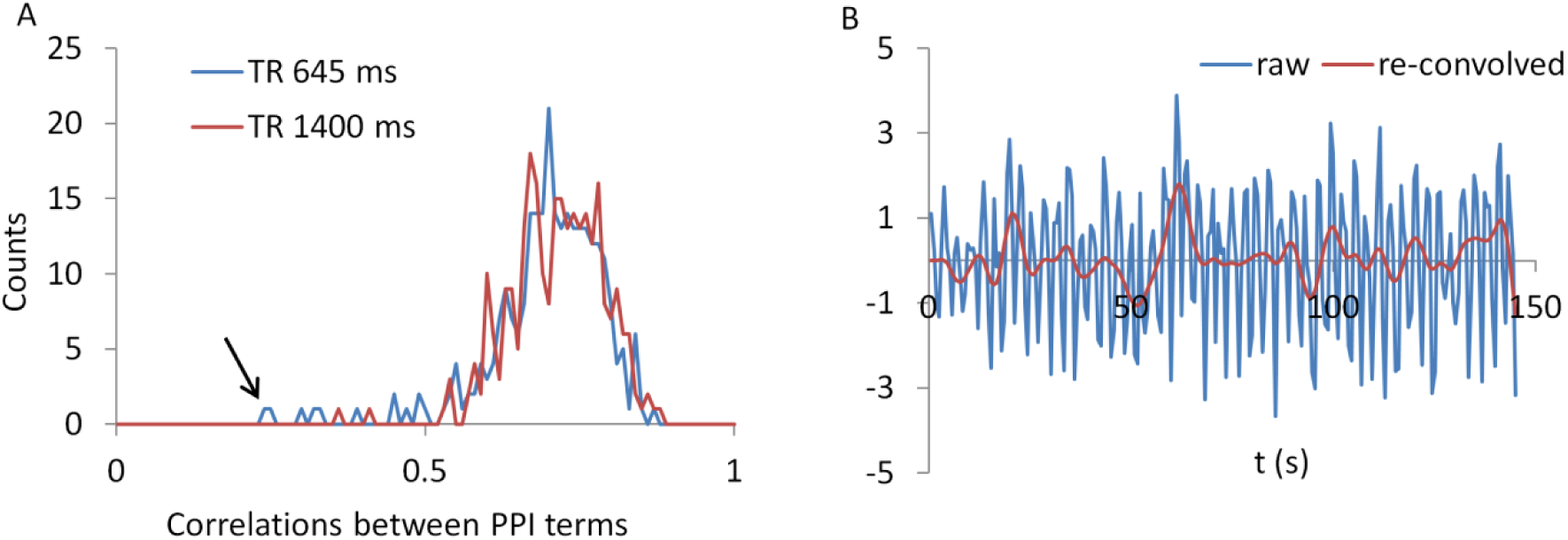
A) Histograms of correlations between PPI terms with and without deconvolution across all subjects from both the LMOG and RMOG ROIs for the two TR runs. B) For the worst case as spotted by the black arrow in A), we show the raw time series and the time series with deconvolution and re-convolution with hemodynamic response function.

We identified the worst case in Figure 12A (black arrow indicated), and deconvolved and reconvovled it with HRF using SPM s method (Figure 12B). The raw and reconvovled signals look dramatically different, with the reconvolved signal resembling a smoothed version of the original signal. Smoothness is indeed the case for the SPM version of deconvolution (Gitelman et al., 2003), because it utilizes regularization to suppress high frequency components of cosine basis functions those were used to approximate the neuronal level physiological variable. To directly illustrate this point, we performed fast Fourier transformation on the time series of the RMOG for all the subjects on the raw, deconvolved, and reconvolved time series for the two TR runs (Figure 13). It could be seen that after deconvolution, high frequency components have been suppressed in both TR runs. Particularly, there is a black line that shows higher power between frequencies of 0.2 to 0.4 Hz in the raw data plot of 645 ms TR run, which coincides to be the outlier observed in Figure 12. The high frequency component was suppressed, so that the reconvolved signal looks smooth.

**Figure 13.**
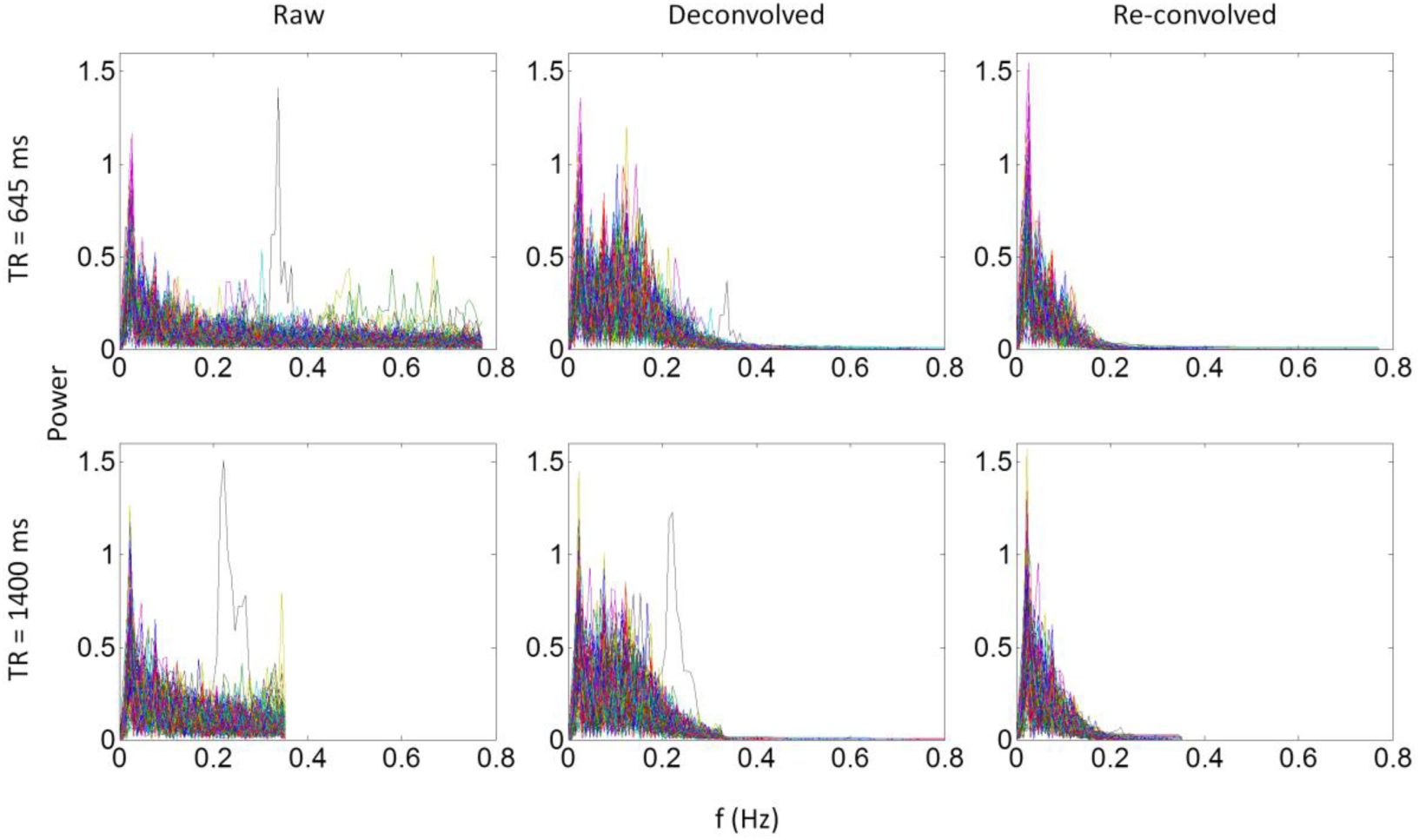
Power spectrums of time series from the right middle occipital gyrus seed for each of the 138 subjects for the 645 ms run (upper panels) and 1400 ms run (lower panels). Each line in a plot represents one subject. Left, middle, and right panels show the power spectrum of the raw, deconvolved, and reconvolved time series, respectively.

## 4. Discussion

By analyzing two separate runs of visual checkerboard task from a large sample (n = 138), the current study first replicated previously reported negative PPI effects between visual cortex and widespread brain regions, and then showed negative PPI effects among visual areas centered in the bilateral fusiform gyrus. By comparing results from two separate runs, we showed that group averaged effects were largely reproducible; however, the inter-subject reliabilities of the PPI effects were typically low. By comparing the deconvolution and non-deconvolution PPI methods, we demonstrated that the results by the two methods were in general very similar, but the non-deconvolution produced larger statistical effects and spatial extents. The non-deconvolution method may reduce inter-subject variations and increase overlaps of results between the two runs in some circumstances compared with the deconvolution method.

### 4.1. Functional connectivity during checkerboard stimulation

The voxel-wise analysis of the LMOG and RMOG seeds replicated our previous results which only analyzed a sub-set of 26 subjects (Di et al., 2017b, 2015). In our previous work (Di et al., 2017b) we could only identify significant PPI effects using the RMOG seed, while the current study demonstrated similar PPI effects from both the LMOG and RMOG seeds. Furthermore, we illustrated that the spatial extent of regions that showed reduced connectivity with the MOG seed could be much larger and extended to other brain regions such as the insula and bilateral sensorimotor cortex. This further suggests a higher extent of functional segregation between the visual cortex and other brain systems during such a simple visual stimulation task compared with the fixation. The current study also extended previous study by analyzing task modulated connectivity effects among cytoarchitectonically defined visual areas. Reduced functional connectivity was observed among many visual areas, with the bilateral FG1 as hub regions. FG1 is the most posterior portion of the fusiform gyrus, which just laid anterior to the occipital cortex (Caspers et al., 2013). It is thought a transition zone between lower retinotopic visual areas and higher category specific brain areas, and integrates information from different retinotopic visual areas to higher category specific brain areas (Caspers et al., 2014). Therefore, it is reasonable to see that the FG1 showed reduced functional connectivity with many lower visual areas in the checkerboard condition, because the simple stimuli cannot form a meaningful percept of a specific category.

The thalamus is a critical subcortical structure in the brain, which not only relay sensory information to the cortex, but also thought to mediate corticocortical communications (Guillery and Sherman, 2002; Saalmann and Kastner, 2011). The PPI analysis of the thalamus, however, did not show consistent effects in different TR runs, and different methods. It may because that the visual thalamus is small in size compared with cortical visual areas, and the signals in the thalamus are not reliable enough. The current results do suggest some reduced connectivity between the visual thalamus to the primary visual cortex, and increased connectivity between the visual thalamus to the anterior portion of the thalamus, basal ganglia, and insula. However, the results are weak and unreliable, especially considering that the current analysis had included 138 subjects.

### 4.2. Reproducibility and reliability of PPI effects

To our knowledge, the current study is the first one to evaluate reproducibility and reliability on PPI effects. The current analysis did not only reproduce the results reported previously (Di et al., 2017b), but also examined the reproducibility between two runs. Although the two runs were scanned using different parameters, most importantly the temporal and spatial resolutions, the patterns of PPI effects turned out to be quite similar between the two runs. The run with 645 ms TR seemed to generate larger spatial extent in the voxel-wise analysis and more statistically significant results in the ROI-wise analysis. This is consistent with our prediction, because there are more time points in the 645 ms TR run than in the 1400 ms TR run, which could yield higher statistical power. We do notice that in some scenarios, i.e. voxel-wise analysis with deconvolution, the PPI results in 645 ms TR run had smaller effect size and spatial extent, which might be due to failure of deconvolution.

On the other hand, the results indicated that inter-subject reliabilities are typically low (around 0.07) no matter which PPI method was used. The low reliability should be compared with those of simple task activations, which showed reasonably high reliability regardless of the scan length. The reliability of PPI effects in the current analysis are also much lower than previous reported test-retest reliabilities on task activations (Plichta et al., 2012; Raemaekers et al., 2007) and resting-state functional connectivity (Guo et al., 2012; Zuo et al., 2010b). Of course the short scan lengths could be one factor that explains the low reliability of PPI effects. But it should be also emphasized that the reliability of higher order interaction effects (i.e. the PPI) should be much lower than the main effects of task activations and task-free functional connectivity. A scan length that is sufficient for obtaining reliable task activations may not be necessarily enough to yield reliable task modulated connectivity estimates. This factor should be taken into account when designing studies on task based connectivity.

### 4.3. Deconvolution and PPI

The PPI results using both the deconvolution and non-deconvolution methods are in general very similar. This is consistent with the simulation showing that the PPI term calculated from the convolution then multiplication method is very similar to the hypothetical PPI term with a known neural activity in a block-designed task. When comparing the differences of PPI results with these two methods, the non-deconvolution method seems to be able to generate larger statistical effects and greater spatial extents or number of significant effects. The non-deconvolution method also increased the Dice coefficients of thresholded PPI maps between the two TR runs. However, the Dice coefficients of thresholded PPI matrices between the two TR runs are quite similar between the two PPI methods, and the deconvolution method may be even benefiting at higher thresholds. These results highlighted the uncertainty of deconvolution method in PPI analysis.

We have shown that the correlations of PPI terms between deconvolution and non-deconvolution methods may have outliers whose correlations were only 0.2 or 0.3 (Figure 12), which is in contrast with the simulation results (Figure 1B). The lower correlations of PPI terms from empirical data compared with the simulations imply that there might be some uncountable variations introduced during the deconvolution/convolution of real fMRI data. Indeed, deconvolution is rather a practical problem to recover underlying signals from some recorded measures, than a simple mathematical problem as depicted in equation 2. In the practical context, measurement noises need to be taken into account in the deconvolution model. For fMRI, the goal of deconvolution is to recover neuronal activities from observed BOLD signals, where there are plenty of noises during MRI recording. The deconvolution should be expressed as follows with an additional error term:

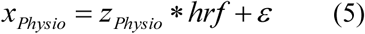

In this circumstance, some noises would be removed during deconvolution so that a signal deconvolved and convolved back with a HRF will no longer be the same as the original signal. The noise characteristics and regularization methods for recovering *zPhysio* become critical to the success of deconvolution.

As have been shown in Figure 13, SPM s deconvolution method explicitly suppresses high frequency components with the intention that the hemodynamic response is slow therefore high frequency components may represent noises. But this may overly smooth the data and remove useful information in higher frequency bands, thus making PPI results with the deconvolution method less sensitive than those with the direct PPI method. This problem may be more severe for short TR data, because there are more high frequency components in the data. On the other hand, high frequency signals in BOLD have been increasingly recognized as functionally meaningful (Chen and Glover, 2015; Gohel and Biswal, 2015; Lewis et al., 2016), and high frequency components may be critical for connectivity dynamics. Given that multiband imaging technique has made fMRI sampling rate much faster, proper treatment of high frequency signals may be critical in deconvolution of fMRI signals and connectivity analysis in general.

Given the facts that the two PPI methods can generate similar results for the current block-designed task and the non-deconvolution method may increase statistical power, we lean toward a conclusion that the non-deconvolution PPI method may be a better choice for a block-designed task. This is in line with the recommendation by FSL (O’Reilly et al., 2012). Of course, deconvolution is still necessary for an event-related task design, because the PPI terms calculated from the convolution then multiplication method are dramatically different from those calculated from the multiplication then convolution method (Figure 1). It s also worth mentioning that it has been suggested that the beta series method (Rissman et al., 2004) might be an alternative method for event-related designed data (Cisler et al., 2014). Lastly, there are indeed many variety of deconvolution methods (Havlicek et al., 2011; Makni et al., 2008; Wu et al., 2013), and some of the methods may be more suitable for fMRI signals and PPI analysis. Systematic comparisons between these different methods are needed in the future.

The current analyses are mostly based on empirical fMRI data. One limitation of empirical analysis is that there is no known ground truth to compare with. Simulation may be an alternative way to approach the question. However, development of biological realistic models for task modulated connectivity is still challenging, so that the deconvolution problem is difficult to study using simulations at the current stage. In addition, the similarities and differences between PPI results of the deconvolution and non-deconvolution methods depend on the variability of hemodynamic response in real fMRI data, which cannot be simply derived from simulations. Therefore, we believe that the current empirical analysis is suitable for the question of deconvolution.

### 4.4. Practical implications on PPI analysis

The current study analyzed data from a simple task design with one task condition and one baseline condition. In real fMRI experiments, however, there are usually more than two conditions. To deal with multiple conditions, it was recommended that each task condition is modeled separately with respect to all other conditions (McLaren et al., 2012). In such “generalized PPI” framework, each experimental condition is modeled as the same way as the checkerboard condition in the current study. It is reasonable to conclude that the similarities of PPI results with and without deconvolution could be generalized to experiments with more than two conditions.

Task related functional connectivity as measured by PPI analysis is typically much smaller, in terms of effect size, reproducibility, and reliability, than simple task activations, and has much larger measurement error. To ensure enough statistical power and reliability, a larger sample size than typical activation studies and enough scan length for each subject are necessary. The design for an fMRI task needs to consider scan length as a critical factor, if the goal of the study is to examine task related connectivity. To date, it is still largely unknown how long a scan is needed for reliability capture task related connectivity. We can only get some insights from resting-state connectivity research, where large scale test-retest datasets are available (Biswal et al., 2010; Zuo et al., 2014). In resting-state literature, it has been suggested that at least five minutes of scan is needed for reliability estimate functional connectivity (Birn et al., 2013; Van Dijk et al., 2010). Then at least five minutes of scan length for a single task condition is needed for task based fMRI. If the PPI effects are going to be compared between two experimental conditions, which is usually the case for a well-designed cognitive neuroimaging study, the required scan length would be much longer. Of course, direct examinations of the effect of scan length on task related connectivity estimates are still needed in future research.

The PPI method takes advantages of the dynamic aspect of the BOLD signals. Therefore, it s preferable to adopt faster sampling rate to capture temporal dynamics, which may in turn lead to sacrifice of other aspects of the signals, e.g. spatial resolution. The current results support the idea that shorter TR may be beneficial for PPI analysis. Of course, faster sampling rate could be accomplished by new developments of MRI techniques such as multi-band acquisition (Feinberg and Yacoub, 2012). However, the current results also suggested some pitfalls of using short TR data. The currently used HRF models and deconvolution method may be not quite suitable for fast TR data, so that the PPI method with deconvolution may fail in some cases in short TR data. More work is still needed to validate and optimize models on high speed fMRI data. Of course, high spatial resolution has its own advantage on mapping small brain structures such as the thalamus. So that the considerations of temporal and spatial resolutions may also need to take into account the spatial scales of regions of interest.

## 5. Conclusion

We demonstrated that the deconvolution and non-deconvolution PPI methods generated similar results on a simple block-designed task. The deconvolution method may be beneficial in terms of statistical power and reproducibility. Taken together, deconvolution may be not necessary for PPI analysis for block-designed fMRI data. When using a large sample, group mean PPI effects are reproducible; however, inter-subject reliabilities of the PPI effects are quite limited. Systematic evaluations on scan length and reliability may be necessary before studying inter-subject differences or group differences of PPI effects.

## Acknowledgement

This research was supported by National Institute of Health grants R01 AG032088 and R01 DA038895.

